# clrDV: A differential variability test for RNA-Seq data based on the skew-normal distribution

**DOI:** 10.1101/2022.09.25.508885

**Authors:** Hongxiang Li, Tsung Fei Khang

## Abstract

Genes that show differential variability between conditions are important for complementing a systems biology understanding of the molecular players involved in a biological process. Under the dominant paradigm for modeling RNA-Seq gene counts using the negative binomial model, tests of differential variability are challenging to develop, owing to dependence of the variance on the mean. The limited availability of methods for detecting genes with differential variability means that researchers often omit differential variability as an analytical step in RNA-Seq data analysis. Here, we describe clrDV, a statistical method for detecting genes that show differential variability between two populations. clrDV is based on a compositional data analysis framework. We present the skew-normal distribution for modeling gene-wise null distribution of centered log-ratio transformation of compositional RNA-seq data. Simulation results show that clrDV has false discovery rate and Type II error that are on par with or superior to existing methodologies. In addition, its run time is faster than the closest competitor’s, and remains relatively constant for increasing sample size per group. Analysis of a large neurodegenerative disease RNA-Seq dataset using clrDV recovers multiple gene candidates that have been reported to be associated with Alzheimer’s disease. Additionally, we find that the majority of genes with differential variability have smaller relative gene expression variance in the Alzheimer’s disease population compared to the control population.

## 1. Introduction

Finding patterns of gene expression variation that are associated with a biological condition of interest is the first step towards elucidating the molecular basis underlying a biological process. Currently, bulk tissue mRNA collected under specific biological conditions through RNA-sequencing (RNA-Seq) technologies remains an important approach for studying gene expression patterns. Typically, genes that show statistically and biologically meaningful difference in mean expression between conditions are of interest. Indeed, pathological conditions frequently manifest as gene sets with altered mean mRNA expression levels. The identification of these genes is important for understanding how the functions of normal molecular pathways are perturbed (Van den Berge *and others*, 2019). Hence, detecting genes that are differentially expressed is a routine and main use of RNA-Seq data (Stark *and others*, 2019). To analyse differential gene expression, a multitude of statistical tests have been developed throughout the years. Methods such as edgeR (Robinson *and others*, 2010), DESeq2 (Love *and others*, 2014) and voom (Law *and others*, 2014) have become established, go-to methods for differential expression (DE) analysis.

To obtain a more complete picture of patterns of gene expression variation, we need to look beyond genes with significantly different mean expression (DE genes) between conditions (Gorlov *and others*, 2012). Genes that show differential variability (DV genes) are likely to be important as well because many biological phenomena are explained by changes in the variance, rather than the mean, of the distribution of gene expression level (de Jong *and others*, 2019). For example, genes that show differential variability between undifferentiated and differentiating states have been found to be related to body axis development, neuronal movement, and transcriptional regulation during the neural differentiation process (Ando *and others*, 2015). In cancer biology, DV genes are useful as biomarkers for predicting tumor progression and prognosis (Dinalankara and Corrada Bravo, 2015), and patient survival (Strbenac *and others*, 2016). Gorlov *and others* (2012) found that genes with larger expression variance in tumors compared to normal cells show stronger association with clinically important features. In network biology, genes with high variability in expression correlate with their positions within the signaling network hierarchy (Komurov and Ram, 2010). Finally, increased gene expression variability is a common outcome of aging (Bahar *and others*, 2006; Stegeman and Weake, 2017). Standard DE analyses are likely to miss DV gene candidates, since they are not optimized for detecting differences in expression variability.

To date, only a few methods are available for finding DV genes using RNA-Seq data. In contrast, even in 2015, there were at least 20 methods for detecting DE genes (Khang and Lau, 2015). For testing differential variability of genes between two populations using RNA-Seq data, initial methods co-opted techniques from microarray data analysis. DiffVar (Phipson and Oshlack, 2014) is an empirical Bayes method that depends on the limma (Smyth, 2005) framework. Subsequently, negative binomial models became popular. MDSeq (Ran and Daye, 2017) uses the coefficient of dispersion (*σ*^2^*/µ*) from the negative binomial 1 (NB1) generalized linear model as a measure for variability. The variance from the NB1 model is a function of the mean *µ* and the dispersion *ϕ* parameter (*σ*^2^ = *ϕµ*). The parameter *µ* is treated as a technical component, whereas *ϕ* is treated as a biological component and interpreted as a parameter for gene expression variability. de Jong *and others* (2019) proposed a DV test that uses the generalized additive models for location, scale and shape (GAMLSS; Rigby and Stasinopoulos (2005)) framework for quantifying expression variability. GAMLSS is based on the negative binomial 2 (NB2) model, whereby the mean and the variance are related quadratically as *σ*^2^ = *µ* + *ϕµ*^2^. Recently, Roberts *and others* (2022) developed DiffDist, a hierarchical Bayesian model based on the NB2 model. In their work, gene expression variability is measured using the dispersion parameter *ϕ*, which is treated as a log-normal prior. Subsequently, test of difference in dispersion between two conditions is based on the posterior distribution simulated using Markov Chain Monte Carlo (MCMC).

In this paper, we wish to propose clrDV - a novel method for detecting DV genes between two conditions in RNA-Seq data that is based on a compositional data analysis framework. The method involves a log-ratio transformation of the raw gene counts, which results in a continuous variable. We show that the skew-normal distribution with centered parameters (Azzalini, 1985) is an appropriate model for the null distribution. Subsequently, we construct a Wald test statistic for testing differential variability. Through simulations, we show how well clr-DV performs compared to existing methods. Finally, we demonstrate the applied value of clrDV by using it to identify biologically meaningful genes in the analysis of a large RNA-seq dataset from a neurodegenerative disease study.

## 2. Motivation

The general idea of conducting a test of differential variability for RNA-Seq data involves testing the equality of variances (equivalently, standard deviations) between two populations. The variance parameter is embedded in some probability distribution that approximates the distribution of gene (more generally, transcript) counts, assuming the null hypothesis is correct. The standard approach models RNA-Seq data as a discrete random variable.

Before modeling can be done, the raw count data need to be normalized to account for variation in the sequencing depth of each sample. Commonly used methods include the trimmed mean of M values (TMM) (Robinson and Oshlack, 2010), the median-of-ratios method (Anders and Huber, 2010; Love *and others*, 2014), upper-quartile (Bullard *and others*, 2010), conditional quartile normalization (Hansen *and others*, 2012), etc. After this, a model that accounts for overdispersion commonly seen in RNA-Seq data (e.g. the NB distribution) is used, but alternative models are possible (Esnaola *and others*, 2013). Statistical tests of differential variability can then be based on estimators of suitable model parameters for representing expression variability.

In recent years, there has been an increasing call towards adopting a compositional data analysis (CoDA) framework for improving the analysis of RNA-Seq data. Indeed, in the closely related field of microbiome data analysis, CoDA forms the main theoretical framework of data analysis and differential abundance methods (Gloor *and others*, 2017). Nevertheless, the diffusion of CoDA approach into RNA-Seq data analysis is slow, possibly because established protocols for routine analyses such as differential expression analysis (e.g. DESEq2, edgeR) are all based on discrete count models such as the NB model. Quinn *and others* (2018) argued that next-generation sequencing abundance data should be viewed as inherently compositional because only a portion of genes may be sampled by sequencers, and cells are likely to be constrained in their capacity for mRNA production. Furthermore, Quinn *and others* (2018) showed the feasibility of applying ALDEx2 (Fernandes *and others*, 2014), a tool developed for differential abundance analysis in micriobiome studies under a CoDA framework, to differential expression analysis using RNA-Seq data. Encouragingly, they reported that ALDEx2 shows superior performance with respect to precision and recall when compared against edgeR and DESeq2. By removing the need to rely on assumptions that justify normalization protocols in standard count-based approaches, log-ratio based transformations of RNA-Seq data in compositional form is potentially more attractive and effective for differential expression analyses (Quinn *and others*, 2019). More recently, McGee *and others* (2019) developed absSimSeq - a novel simulation protocol for generating realistic RNA-Seq data using a compositional data framework.

The key step in processing compositional data involves a log-ratio transformation, for which several variants are available. The simplest is the centered log-ratio (CLR) transformation, first proposed by Aitchison (1986). After CLR-transformation, the simplex space of the compositional data is transformed into the Euclidean space. It is then convenient to view CLR-transformed values as realizations of a continuous random variable. To be concrete, let *X*_*gi*_ be the read count for gene *g* and sample *i*, where *g* = 1, 2, …, *G* and *i* = 1, 2, …, *n*. For a *G*-component composition {*x*_1*i*_, *x*_2*i*_, …, *x*_*Gi*_}, the CLR-transformation of *X*_*gi*_ is given by

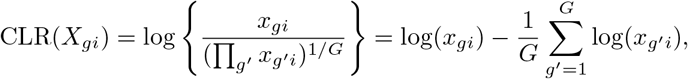

for *g*^*t*^ = 1, 2, …, *G*. We call CLR(*X*_*gi*_) the relative gene expression, or CLR-transformed count, of gene *g* and sample *i*. A pseudo-value 0.5 is added if *x*_*gi*_ = 0 for any *i*. Thus, the main challenge for using CLR-transformed data to develop a test for differential variability is modeling them using a tractable probability distribution for which estimation of the variance parameter is practical.

## 3. Materials and Methods

### 3.1 The skew-normal model for CLR-transformed data

We show that the null distribution of CLR-transformed count data approximately follows the skew-normal distribution (Azzalini, 1985; Azzalini and Capitanio, 2014; see Supplementary Material S1). Denote the relative gene expression from gene *g* in sample *i* by *Y*_*gi*_. Thus, *Y*_*gi*_ has a skew-normal distribution with centered parameters (CP), that is, *Y*_*gi*_ ∼ SN_C_(*µ*_*g*_, *σ*_*g*_, *γ*_*g*_), where *µ*_*g*_ is the mean, *σ*_*g*_ is the standard deviation, and *γ*_*g*_ is the skewness parameter, *g* = 1, 2, …, *G* and *i* = 1, 2, …, *n*. The parameter vector 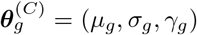 has parameter space ℝ × ℝ^+^ × (−*k, k*), where 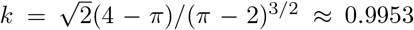. The special case of *γ*_*g*_ = 0 results in a normal distribution with mean *µ*_*g*_ and variance 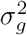. The probability density function of a skew-normal distribution with direct parameters (DP) is given by

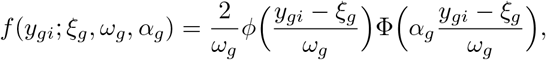

with location parameter *ξ*_*g*_ ∈ ℝ, scale parameter *ω*_*g*_ ∈ ℝ+, and skewness parameter *α*_*g*_ ∈ ℝ; *ϕ*(·) and Φ(·) are the probability density function and the cumulative distribution function of the standard normal distribution, respectively. The skew-normal distribution with CP is derived from the DP form via the mapping (Azzalini and Capitanio, 2014)

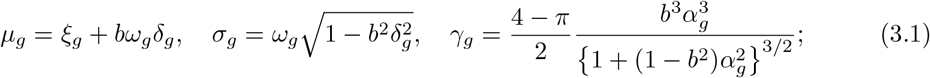

and the inverse mapping is provided by

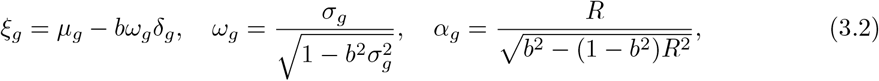

where 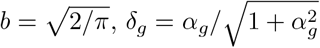, and 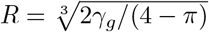.

For a single sample, the log-likelihood function for 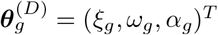 is given by

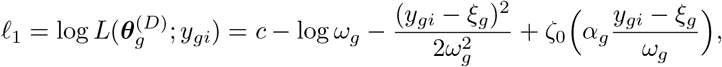

where *c* is a constant and *ζ*_0_(·) = log {2Φ(·)}. Taking *z*_*gi*_ = (*y*_*gi*_ − *ξ*_*g*_)*/ω*_*g*_, we obtain the partial derivatives of *l*_1_:

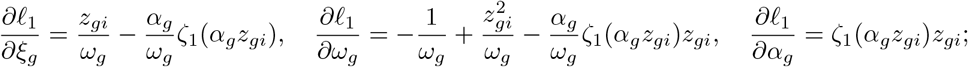

thus the likelihood equations for a sample of size *n* are given by

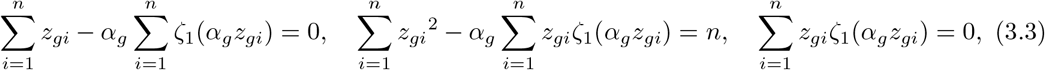

where *ζ*_1_(·) = *ϕ*(·)*/*Φ(·). Numerical methods are necessary to solve these equations. Azzalini and Capitanio (2014) suggested that a sample size up to about 50 may be necessary for the skew-normal distribution. To initialize the search, method of moments (MM) estimates are chosen as starting points for the CP components in Eq. (3.1). The MM estimators for the centered parameters are given by

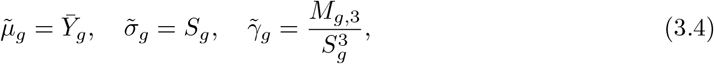

respectively, where 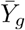 is the sample mean, *S*_*g*_ is the sample standard deviation, and *M*_*g*,3_ is the sample third central moment. By estimating the CP components in Eq. (3.1) using Eq. (3.4), and then converting them to DP components using Eq. (3.2), we obtain the MM estimators of the DP components: 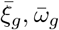 and 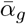. Subsequently, a search of the DP space where Eq. (3.3) holds is done. Once 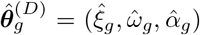 is obtained, it is mapped to Eq. (3.1) to get 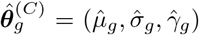, the maximum likelihood estimators of the centered parameters.

Under regular maximum likelihood estimation, certain data values can produce a divergent 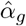. To overcome this problem, Azzalini and Arellano-Valle (2013) proposed a maximum penalized likelihood estimation (“Qpenalty”) approach. A non-negative penalty term *Q* that penalizes the divergence of the skewness parameter *α*_*g*_ is formulated as 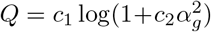, where *c*_1_ ≈ 0.87591 and *c*_2_ ≈ 0.85625 (Azzalini and Arellano-Valle, 2013; Azzalini and Capitanio, 2014). Then, the maximum penalized likelihood for 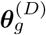 is the penalized log-likelihood

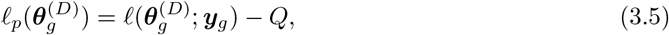

where 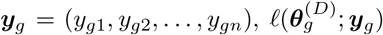 is the log-likelihood function with respect to the parameter vector 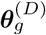:

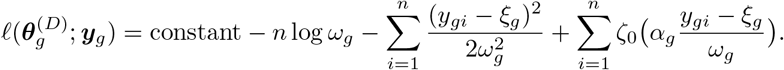

The maximum penalized likelihood estimator (MPLE), 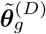, is a finite point that maximizes 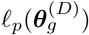. The standard errors of 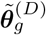 can be approximated from the corresponding penalized information matrix as Var 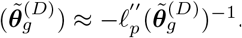.

The “MPpenalty” approach (Azzalini and Capitanio, 2014) defines the penalty function *Q* in Eq. (3.5) as − log *π*_*m*_(*α*_*g*_), where *π*_*m*_ is a prior distribution for the skewness parameter *α*_*g*_. The matching prior (Cabras *and others*, 2012) for *α*_*g*_, allowing for the presence of ***ψ*** = (*ξ*_*g*_, *ω*_*g*_), is given by

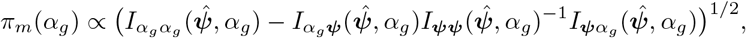

where the terms involved are specific blocks of the Fisher information matrix ***I*** of 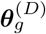 (see Supplementary Material S1 for details). Since *π*_*m*_(0) = 0, the matching prior penalty effectively penalizes *α*_*g*_ = 0 with *Q* = ∞.

To perform parameter estimation and carry out related numerical tasks involving the skew-normal distribution, we used the `sn` R package (Azzalini, 2022). Regular maximum likelihood estimation of parameters of the skew-normal model was first done using the function `selm()`. If NA values were returned, we used the maximum penalized likelihood estimation as implemented using the `Qpenalty` option. If NA values persisted, the `MPpenalty` option was used.

For RNA-Seq experiments comparing two populations, testing for differential variability is equivalent to testing the equality of the standard deviation of relative gene expressions in two populations, that is, *σ*_*g*,1_ = *σ*_*g*,2_. For this purpose, we can use the Wald statistic

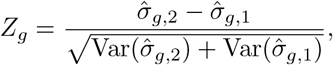

for *g* = 1, 2, …, *G*, where 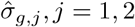 are the maximum likelihood estimators of the standard deviation of the skew-normal distribution with centered parameters for population 1 and population 2, and Var 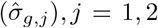 are diagonals of the estimated Fisher information matrix of centered parameters 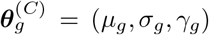. To control the false discovery rate (FDR) as a result of conducting multiple independent hypothesis tests across genes, we applied the Benjamini-Yekutieli (Benjamini and Yekutieli, 2001) procedure.

### 3.2 Data Description

In order to study the performance of clrDV and other existing methods with respect to FDR and Type II error, it is necessary to simulate the null distribution with realistic parameter values. For this purpose, we used two real RNA-Seq datasets. The first dataset (GEO accesion number: GSE123658) contains whole blood RNA-Seq data from from 39 Type 1 diabetes patients and 43 healthy donors (Leal Valentim *and others*, 2020), with 16, 785 transcripts. The second dataset (GEO acccesion number: GSE150318) contains longitudinal gene expression data from 114 short-lived killfish *Nothobranchius furzeri* measured at 10 weeks and 20 weeks of age (Kelmer Sacramento *and others*, 2020), with 26, 739 transcripts. Hereafter, we call these two datasets the “Valentim dataset” and the “Kelmer dataset”.

For empirical assessment, we used the Mayo RNASeq dataset (Allen *and others*, 2016,?; available at AMP-AD (2022)), which consists of 278 samples and 64, 253 transcripts. RNA was isolated from the temporal cortex of brains of patients with four biological conditions: control (*n* = 80), Alzheimer’s disease (AD; *n* = 84), progressive supranuclear palsy (PSP; *n* = 84) and pathologic aging (*n* = 30). We chose to compare the control group against the AD and the PSP group respectively, since the sample sizes in these groups are reasonably large and balanced.

### 3.3 Simulation study

Only transcripts that satisfy two conditions in each group were used for simulation: (i) average count-per-million (CPM) above 0.5; and (ii) less than 85% of samples have zero count. Then, 2000 of the filtered genes were randomly selected. For each gene, an NB2 model was fitted. We simulated 10% of the genes to be DV genes by multiplying their size parameter (1*/ϕ*) with a random value *x*, where *x* ∈ (0.25, 0.5)∪(2, 4). Counts were then simulated based on the fitted NB2 model, for six sample sizes (50, 100, 125, 150, 200, 250) using the `polyester` R package (Frazee *and others*, 2015). A total of 30 instances were thus simulated. Genes with BY-adjusted *p*-value *<* 0.05 were flagged as having differential variability.

The performance of clrDV against MDSeq, diffVar, and GAMLSS (Benjamini-Hochberg (BH) and Benjamini-Yekutieli (BY) variants) was evaluated by considering their FDR and Type II error. Additionally, we also recorded the run time of each method. DiffDist was excluded from the evaluation since it needs to perform MCMC simulations to generate the posterior distribution. As such, it is computationally expensive to implement and difficult to justify as a choice for routine application. Indeed, running DiffDist on an RNA-Seq dataset with 43 samples per group and 23, 416 transcripts, Roberts *and others* (2022) reported that DiffDist took about three hours to complete, compared to 12 minutes for GAMLSS and 4 minutes for MDSeq.

### 3.4 Empirical assessment

We applied clrDV to the Mayo RNA-Seq dataset to assess its capacity for detecting DV genes that are contextually meaningful. Analysis using MDSeq and GAMLSS (BH and and BY variants) were also done. We dropped diffVar because this method performed poorly during the simulation stage. Volcano plots were used to inspect the biological effect size and statistical significance of all genes tested. Venn diagrams were used to identify sets of genes that are identically recovered by all three methods, by combinations of two methods, or uniquely recovered by a single method. Violin plots of selected DV genes were made to verify computational results.

### 3.5 Tools and computing environment

Computational tasks were done in a computer with a 1.80 GHz i5-8265U CPU and an 8GB RAM processor. R (version 4.2.1) (R Core Team, 2022) operating in Windows 10 was used. The complete list of R packages used is given in the Supplementary Material S2. ENSEMBL gene ID to gene symbol conversion was done using the application programming interface of the BioTools.fr website (Saurin, 2022).

## 4. Results

### 4.1 Simulation study

We found that the skew-normal distribution with centered parameters fit the CLR-transformed count data well. Two examples are given in Figure 1. Figure 2 shows the scatter plots of Type II error against FDR for analysis of the simulated Valentim dataset, for each of the six sample size per group scenarios. diffVar is clearly uniformly inferior to all other methods (mean Type II error *>* 0.05 and FDR *>* 0.17, for all sample sizes). For sample size of 50, all methods show relatively larger mean Type II error (*>* 0.2); additionally, diffVar and GAMLSS-BH show high mean FDR (*>* 0.05). Against MDSeq, clrDV is uniformly superior with respect to mean FDR and mean Type II error; against GAMLSS-BH, clrDV has uniformly superior mean FDR; against GAMLSS-BY, clrDV gives approximately similar mean FDR and mean Type II error. When sample size is very large (250), clrDV, MDSeq and GAMLSS-BY give similar performance. With respect to computing speed, clrDV is substantially faster than GAMLSS (both BH and BY variants) as sample size increases (Table 1). For the analysis of simulated data from the Kelmer dataset, we find clrDV to have comparable mean FDR and mean Type II error (Fig. 3) as MDSeq and GAMLSS-BY. However, clrDV computing time remains almost constant across the six sample sizes, whereas MDSeq and GAMLSS have computing times that increase with sample size (Table 2). diffVar and GAMLSS-BH are inferior in controlling FDR across all six sample sizes.

**Fig. 1.**
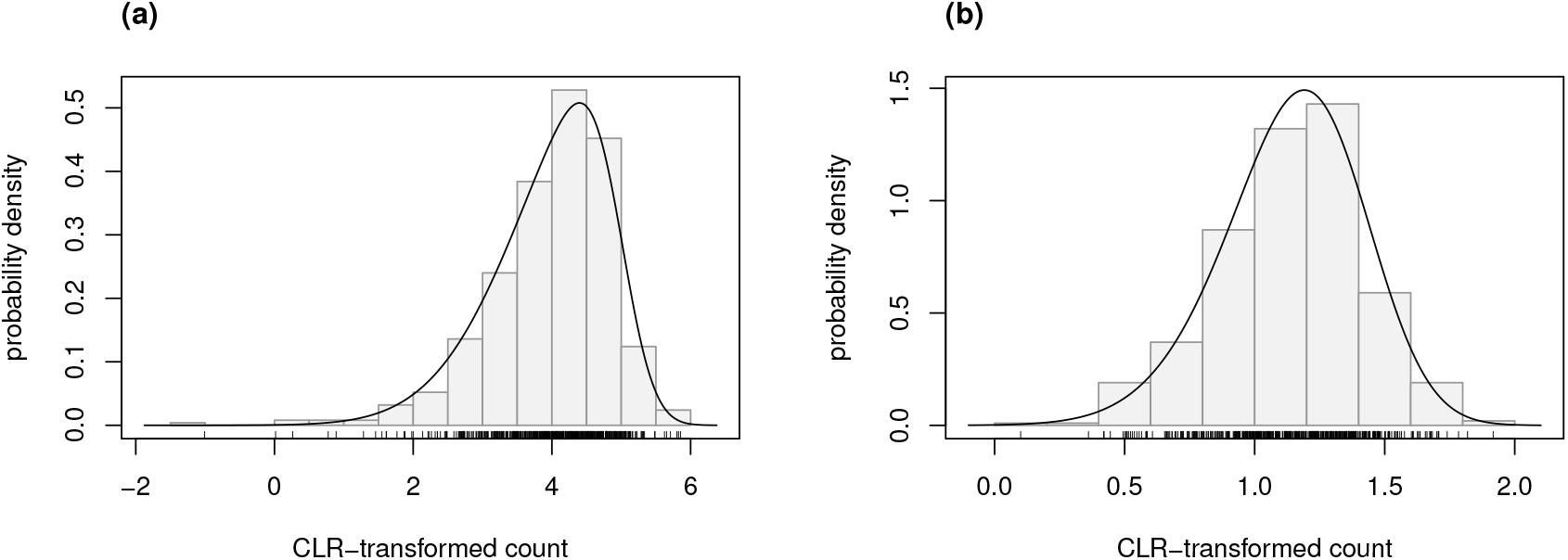
Histograms of CLR-transformed counts for two genes with fitted skew-normal curve for (a) the Valentim dataset 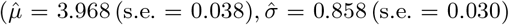 and 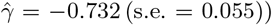 (b) the Kelmer dataset 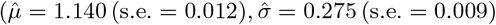 and 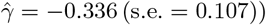.

**Fig. 2.**
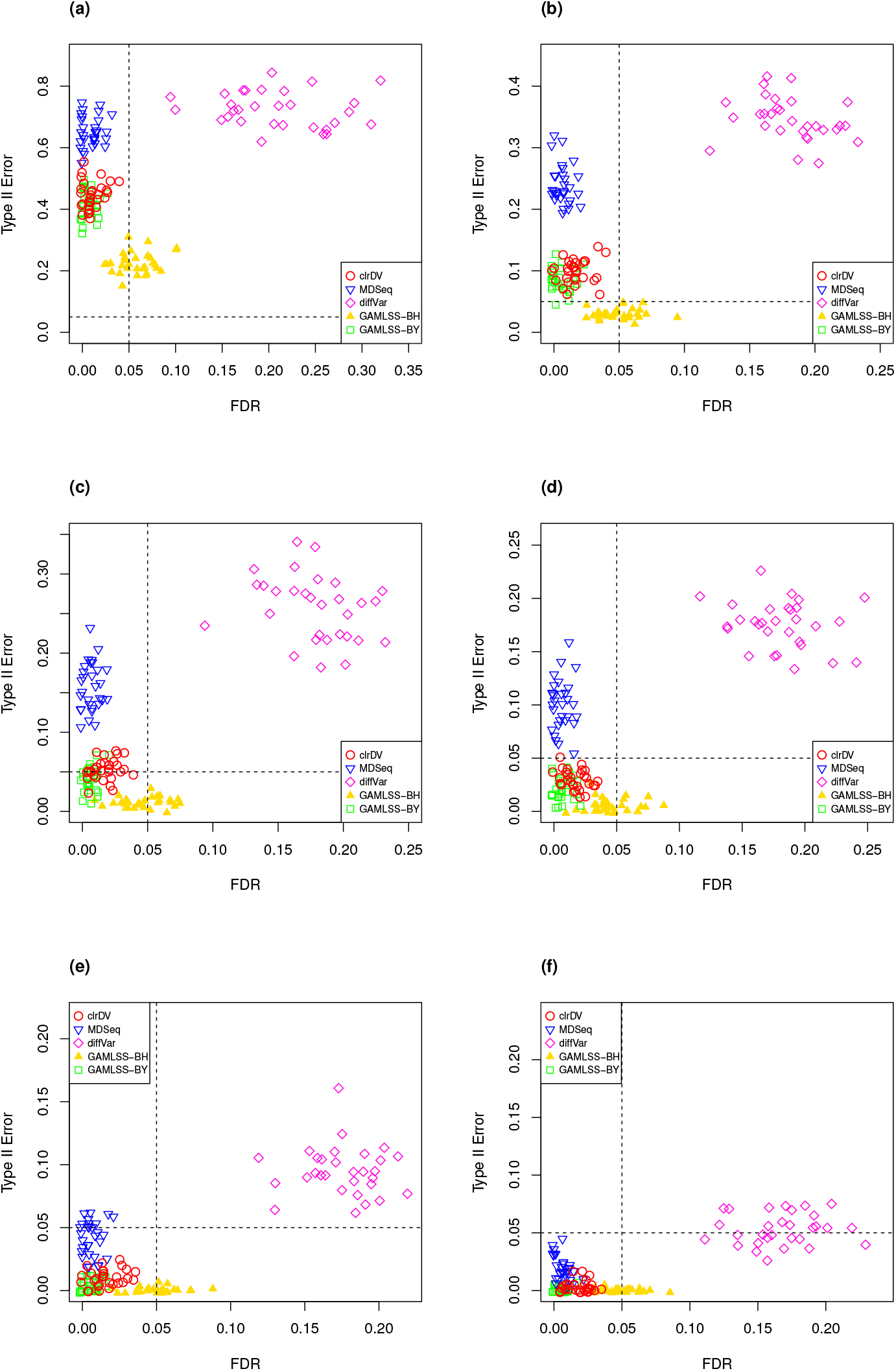
Scatter plots of Type II error vs. FDR for simulation study of the Valentim dataset (30 instances) for samples size per group of (a) 50, (b) 100, (c) 125, (d) 150, (e) 200, and (f) 250. Dashed lines represent Type II error and FDR of 0.05.

**Table 1:** Mean computing times (in seconds) for each of the four DV tests applied to data simulated from the Valentim dataset (30 instances). Standard deviation in parentheses. BH and BY variants of GAMLSS have similar computing time.

| Sample size<br>per group | Method |  |  |  |
| --- | --- | --- | --- | --- |
|  | clrDV | MDSeq | diffVar | GAMLSS |
| 50 | 81(5) | 59(2) | 0.2(0.01) | 79(1) |
| 100 | 69(11) | 103(10) | 0.4(0.2) | 97(12) |
| 125 | 71(9) | 126(10) | 0.6(0.2) | 106(6) |
| 150 | 67(2) | 143(7) | 0.7(0.3) | 116(1) |
| 200 | 76(8) | 196(14) | 1.3(0.4) | 153(16) |
| 250 | 81(8) | 235(17) | 1.2(0.3) | 170(17) |

**Fig. 3.**
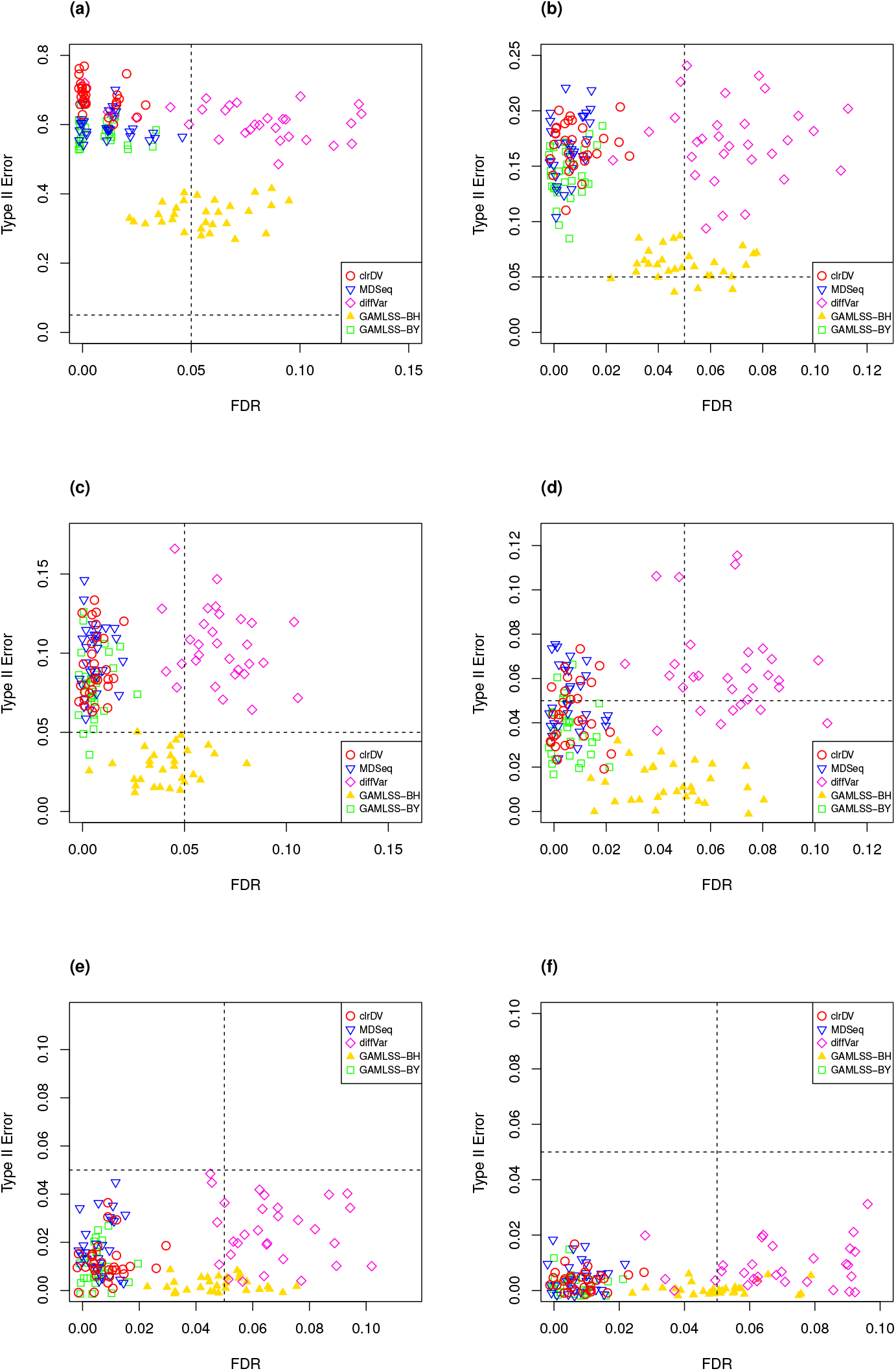
Scatter plots of Type II error vs. FDR for simulation study of the Kelmer dataset (30 instances) for samples size per group of (a) 50, (b) 100, (c) 125, (d) 150, (e) 200, and (f) 250. Dashed lines represent Type II error and FDR of 0.05.

**Table 2:** Mean computing times (in seconds) for each of the four DV tests applied to data simulated from the Kelmer dataset (30 instances). Standard deviation in parentheses. BH and BY variants of GAMLSS have similar computing time.

| Sample size<br>per group | Method |  |  |  |
| --- | --- | --- | --- | --- |
|  | clrDV | MDSeq | diffVar | GAMLSS |
| 50 | 73(5) | 32(1) | 0.2(0.02) | 62(1) |
| 100 | 53(2) | 45(2) | 0.4(0.1) | 70(3) |
| 125 | 58(1) | 57(3) | 0.5(0.2) | 76(0) |
| 150 | 58(2) | 63(3) | 0.7(0.3) | 81(2) |
| 200 | 65(1) | 84(4) | 1.0(0.3) | 95(1) |
| 250 | 71(7) | 104(8) | 1.4(0.2) | 114(6) |

### 4.2 Analysis of the Mayo RNA-Seq dataset

After filtering, sample sizes of the control, the AD and the PSP groups were 78, 82, and 84, respectively. For the AD and the control group comparison, a total of 18,664 transcripts were left; for the PSP and control comparison, 18,636 transcripts were left. For MDSeq and GAMLSS, we normalized the raw counts using TMM normalization.

#### 4.2.1 Detection of genes with differential variability

Applying the procedure described in Section 3, we estimated the standard deviation of the CLR-transformed data, computed the Wald statistic and subsequently the BY-adjusted p-value for each tested gene. For the control vs. AD comparison, we detected a set of 4754 DV genes (see Supplementary Table 1 for complete list); for the control vs. PSP comparison, 4859 DV genes were detected (see Supplementary Table 2 for complete list). For the majority of DV genes, the estimated standard deviation in the control group is larger than the one in the treatment group (Fig. 4). This observation suggests that genes with decreased expression variability among patients with AD are far more common than those that show increased variability.

**Fig. 4.**
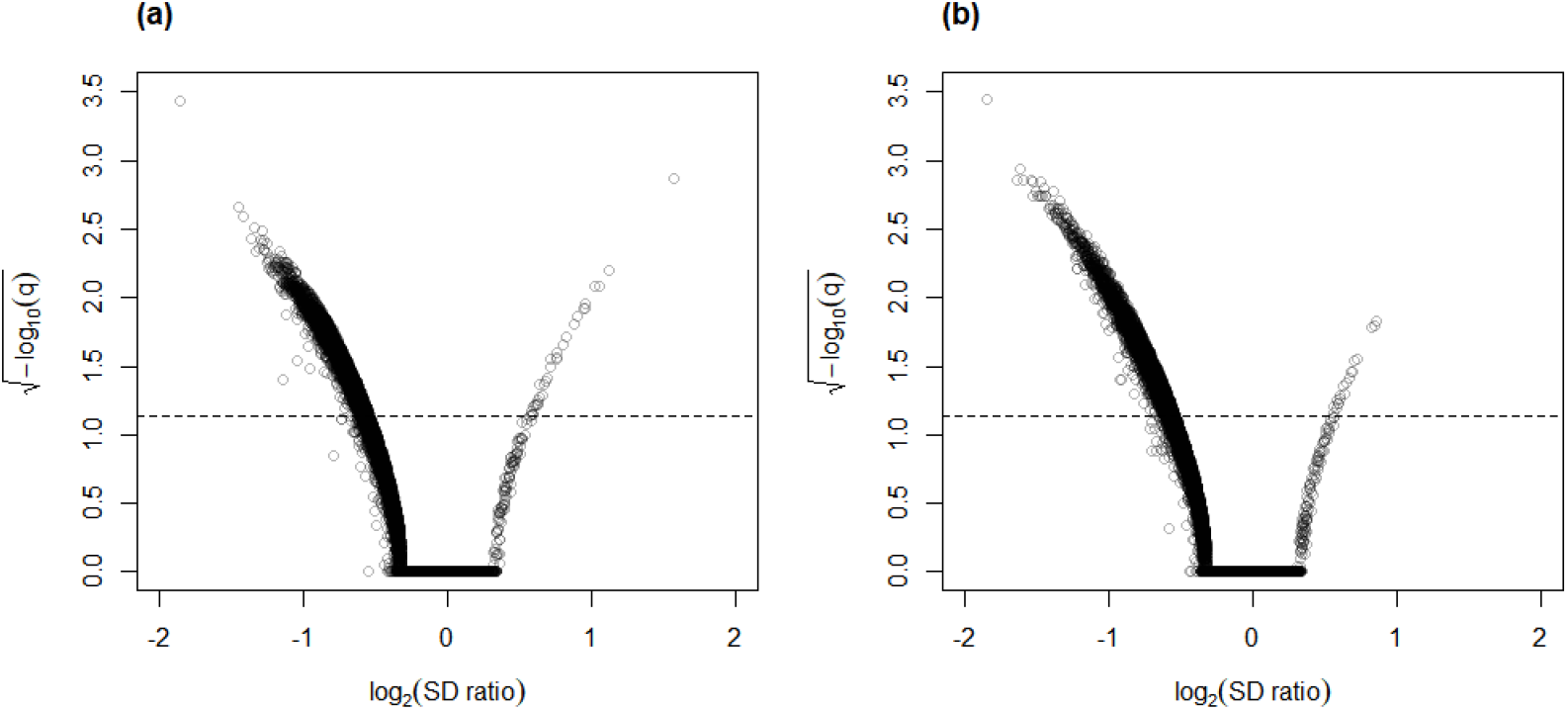
Volcano plots for (a) control vs. AD and (b) control vs. PSP comparisons for the Mayo RNA-Seq dataset. Dashed line represents the threshold of BY-adjusted p-value (*q*) at 0.05 for flagging DV genes. The number of DV genes with log_2_(SD ratio) *>* 0 and log_2_(SD ratio) *<* 0 respectively: (a) 32 and 4722; (b) 19 and 4840.

Figure 5 shows the number of significant DV genes identified by clrDV, MDSeq, GAMLSS-BH and GAMLSS-BY for the control vs. AD comparison (see Supplementary Table 3 for complete list). GAMLSS-BH detected the most DV genes (9926), followed by MDSeq (6944), and clrDV (4754). The high confidence gene set, defined as the intersection of DV genes from each method, contains genes with estimated log_2_(SD ratio) that is relatively large (*>* 0.5). About 99.8% (4745*/*4754) of DV genes detected by clrDV are also identified by MDSeq or GAMLSS-BH; 92.5% (4396*/*4754) are detected by both MDSeq and GAMLSS-BH; about 0.2% (9*/*4754) are uniquely identified by clrDV. GAMLSS-BH identified very large numbers of DV genes in this dataset, but the majority of these are probably false positives, given its relatively poorer control of FDR as shown in the results of the simulation studies. Moreover, these DV genes have estimated log_2_(SD ratio) with relatively small magnitude, as indicated by the violin plots (Fig. 5(c)).

**Fig. 5.**
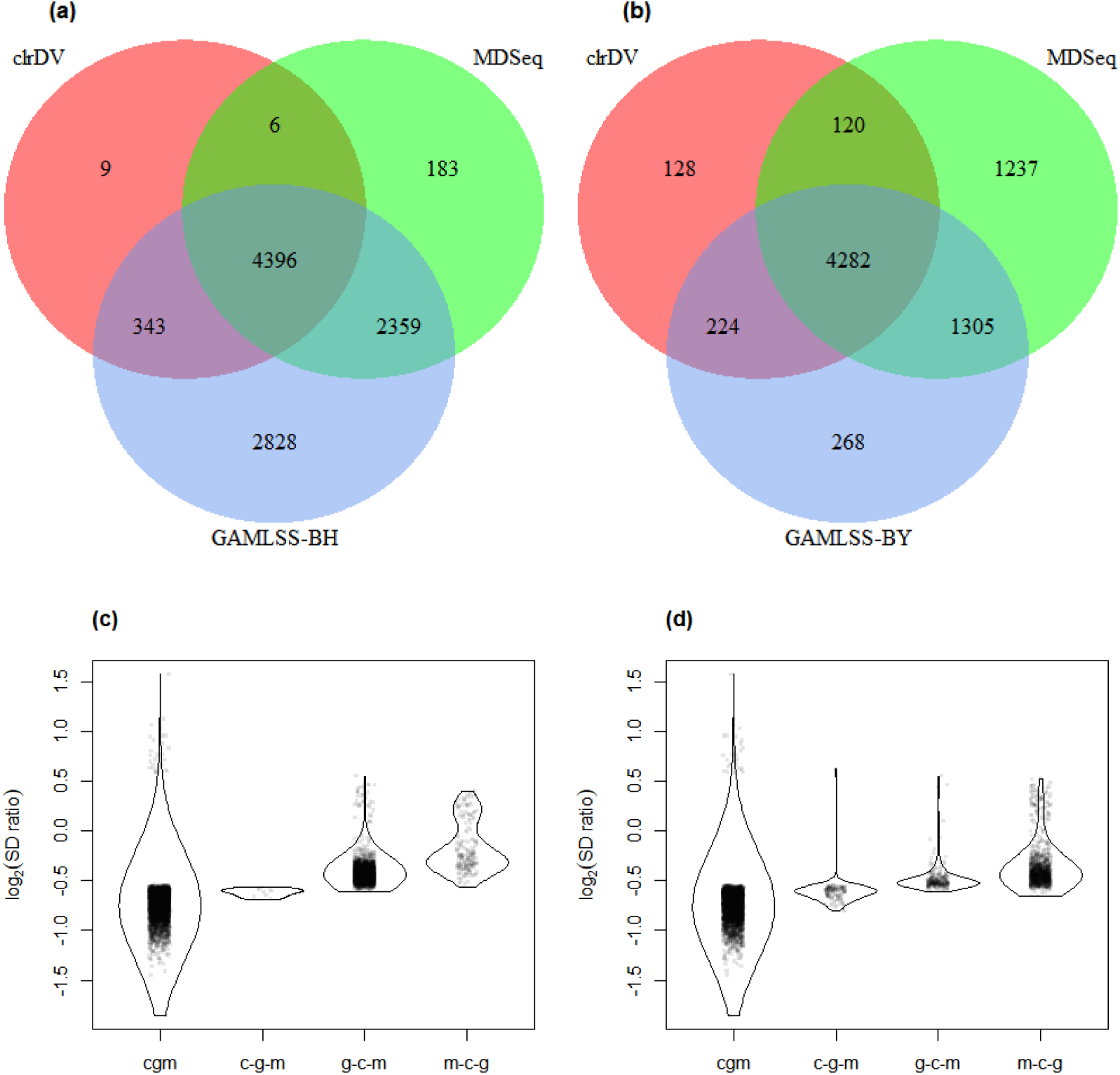
Venn diagrams of DV genes detected by clrDV, MDSeq and (a) GAMLSS-BH; (b) GAMLSS-BY for the control vs. AD comparison. Violin plots of the distribution of estimated log_2_(SD ratio) of the DV genes detected using clrDV, MDSeq and (c) GAMLSS-BH; (d) GAMLSS-BY. Abbreviations: cgm = DV genes detected by clrDV, GAMLSS and MDSeq; c-g-m = DV genes detected by clrDV only; g-c-m = DV genes detected by GAMLSS-BH only; m-c-g = DV genes detected by MDSeq only.

Using GAMLSS-BY, only 6079 DV genes were detected, compared to 9926 using GAMLSS-BH. Thus, GAMLSS-BY primarily helps improve FDR by reducing the number of DV genes called. Between 61.7% (4282*/*6944) and 90.1% (4282*/*4754) of the DV genes detected by one method are detected by all three. About 97.3% (4626*/*4754) of DV genes detected by clrDV are identified by one of other two methods, and 2.7% (128*/*4754) of DV genes detected by clrDV are unique.

The result of the control vs. PSP comparison is similar (Fig. 6; Supplementary Table 4). GAMLSS-BH also detected the most number of DV genes (9707), followed by MDSeq (6920), and clrDV (4859). Up to 99.4% (4831*/*4859) of DV genes identified by clrDV are detected by MDSeq or GAMLSS-BH; about 89.8% (4363*/*4859) are detected by both MDSeq and GAMLSS-BH; about 0.6% (28*/*4859) are uniquely identified by clrDV. Using GAMLSS-BY, only 6024 DV genes were flagged. Approximately 96.1% (4671*/*4859) of DV genes identified by clrDV are also identified by MDSeq or GAMLSS-BY; about 86.6% (4207*/*4859) are detected by both MDSeq and GAMLSS-BY; about 3.9% (188*/*4859) are uniquely detected by clrDV.

**Fig. 6.**
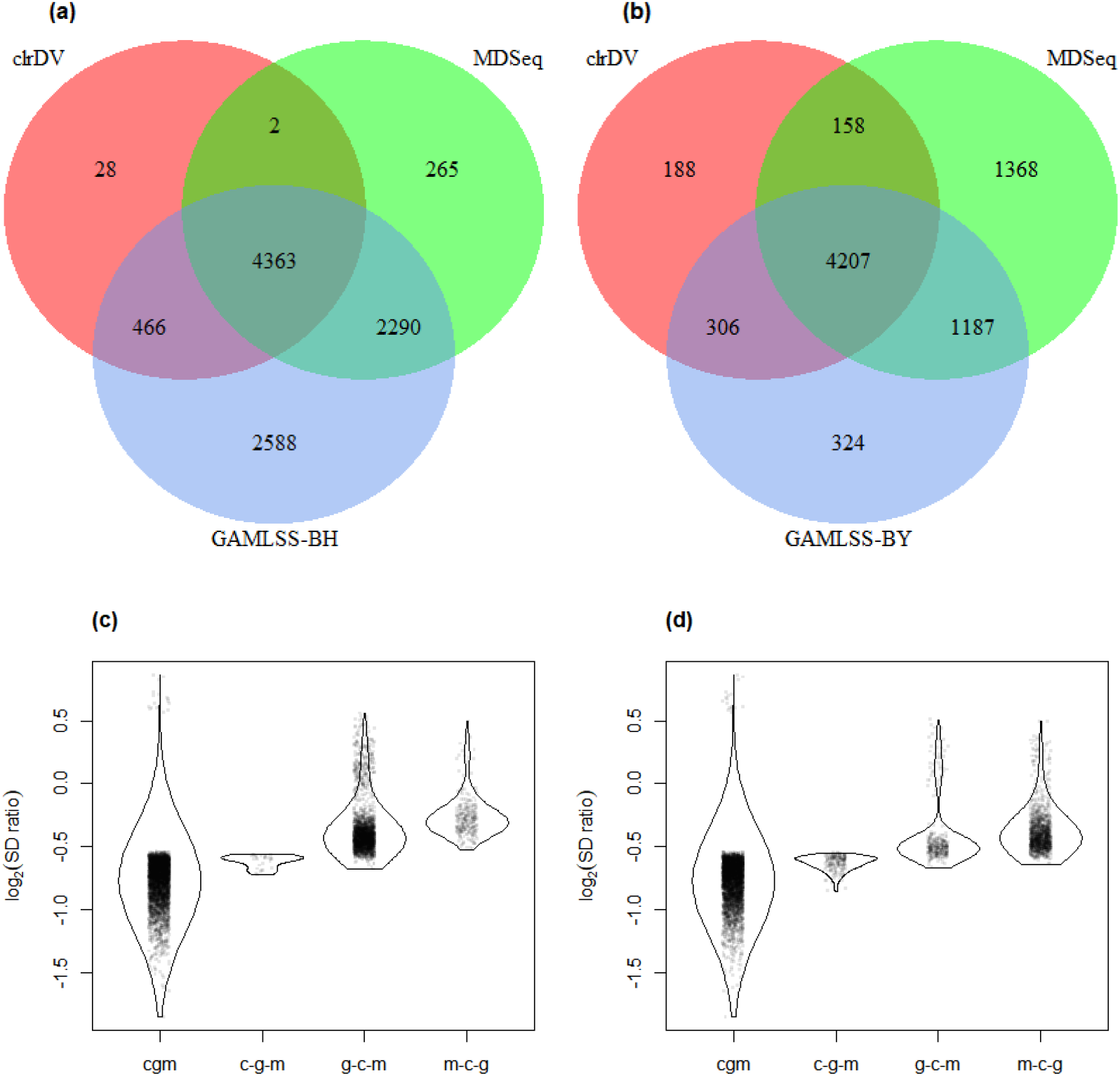
Venn diagrams of DV genes detected by clrDV, MDSeq and (a) GAMLSS-BH; (b) GAMLSS-BY for the control vs. PSP comparison. Violin plots of the distribution of estimated log_2_(SD ratio) of the DV genes detected using clrDV, MDSeq and (c) GAMLSS-BH; (d) GAMLSS-BY. Abbreviations: cgm = DV genes detected by clrDV, GAMLSS and MDSeq; c-g-m = DV genes detected by clrDV only; g-c-m = DV genes detected by GAMLSS-BH only; m-c-g = DV genes detected by MDSeq only.

**Fig. 7.**
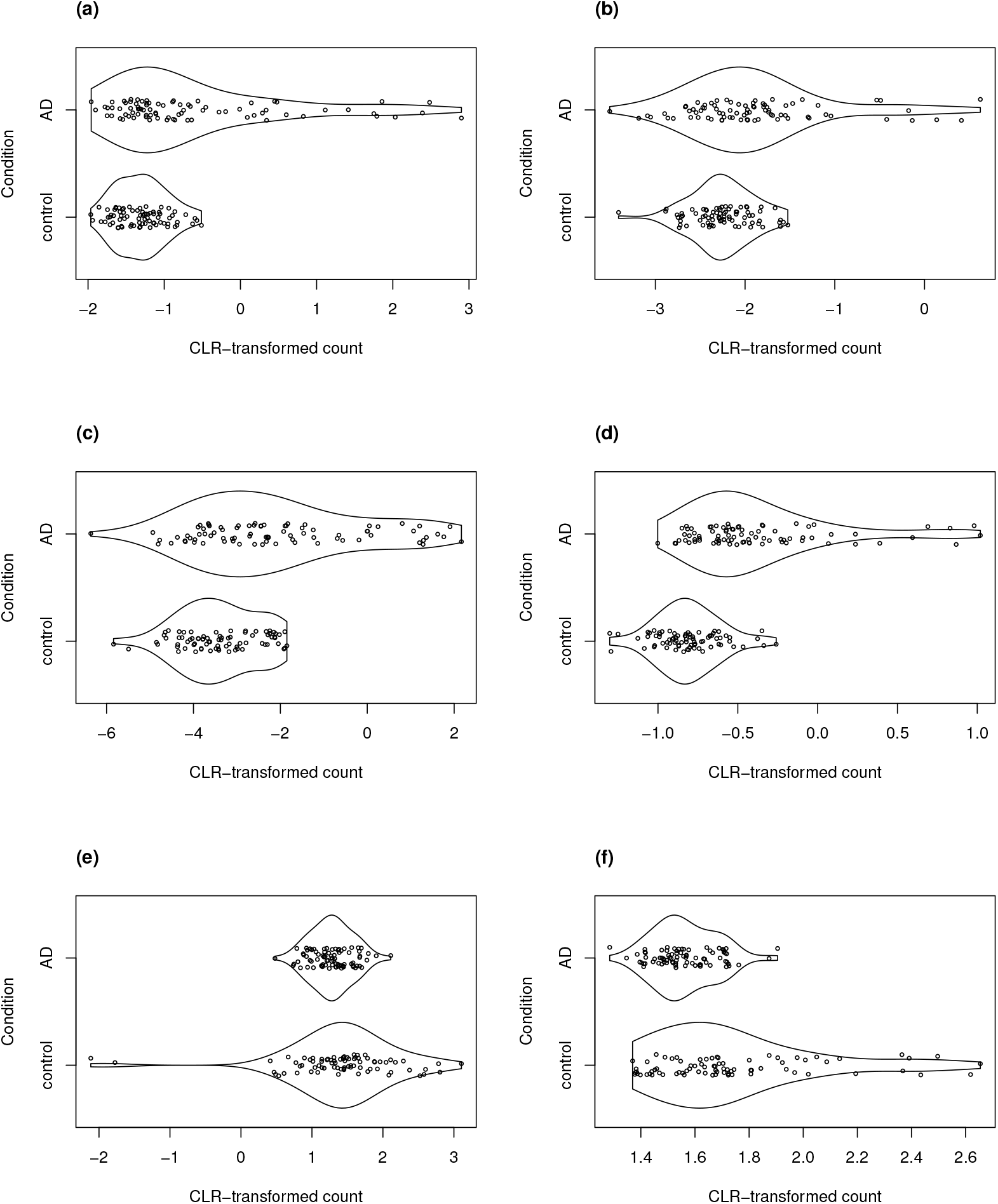
Violin plots of selected DV genes detected in the control vs. AD comparison. (a) SLC47A1, (b) C2orf40, (c) SLPI, (d) LTBP2 have the largest SD ratio (*>* 2); (e) GP1BB and (f) PELP1 have SD ratio about 0.4.

The violin plots (Fig.5 and Fig.6) suggest that the DV genes uniquely called by clrDV may be more likely to true positives, given that the magnitude of log_2_(SD ratio) is generally larger than 0.5. For those uniquely called by GAMLSS or MDSeq, the order of magnitude is generally below 0.5. With respect to run time, for the control vs. AD comparison, clrDV took about 7.5 minutes, compared to 6 minutes for MDSeq, and 13 minutes for GAMLSS; for the control vs. PSP comparison, clrDV took about 7 minutes, while MDSeq used 6 minutes, and GAMLSS used 15 minutes.

#### 4.2.2 Biological significance of detected differential variability genes

In the control vs. AD comparison, four of the DV genes that have the largest estimated SD ratio above 1 are LTBP2, SLPI, C2orf40, and SLC47A1. All four genes have been reported to be associated with Alzheimer’s disease in the literature. The latent transforming growth factor (TGF)-beta binding proteins (LTBP) are important components of the extracellular matrix (Robertson *and others*, 2015). They interact with fibrillin microfibrils, and are known to be mediators of TGF-*β* functions (Rifkin *and others*, 2018), dysfunctions of which have been implicated in Alzheimer’s disease (Das and Golde, 2006). Then, the secretory leukocyte protease inhibitor protein (SLPI) is known to regulate the penetrance of frontotemporal lobar degeneration (FTLD) in patients who have mutations in the progranulin gene (Ghidoni *and others*, 2014). Loss of progranulin function has been found to enhance microglial neuroinflammation, which is implicated in Alzheimer’s disease (Mendsaikhan *and others*, 2019). Podvin *and others* (2016) found that C2orf40 is a neuroimmune factor in Alzheimer’s disease. The SLC47A1 (solute carrier family 47 member 1) protein is expressed in both the kidney and the brain, and recent research has suggested a linkage between kidney diseases and Alzheimer’s disease (Shi *and others*, 2018; Kelly and Rothwell, 2022).

We detected 74 genes from the SLC family in the high confidence DV gene set, including four members of the SLC39 family. Lang *and others* (2012) demonstrated the modulating effect of dZip1, the ortholog of human SLC39 family transporter, on zinc ion uptake using a *Drosophila* model. Zinc is known to induce amyloid beta formation (Bush *and others*, 1994). Inhibition of dZip1 produces substantial reduction of amyloid beta peptide 42 (A*β*42) fibril deposits and less neurodegeneration in A*β*42-transgenic flies.

Two of the DV genes with estimated SD ratio substantially smaller than 1 are PELP1 and GP1BB. PELP1 mediates E2 inhibition of GSK3*β*, a neurodegenerative kinase signaling pathway in the brain (Thakkar *and others*, 2018). GSK3*β* is implicated in Alzheimer’s disease as a key mediator of cell death (Llorens-Martin *and others*, 2014). The GP1BB gene produces glycoprotein 1b-beta (GPIb*β*), a subunit of the GPIb-IX-V protein complex on the surface of platelet cells. Amyloid beta peptides are known to be actively released by platelets (Bush *and others*, 1990; Casoli *and others*, 2007). Visconte *and others* (2020) recently reported that recruitment of GPIb-IX-V is required for fibrillar amyloid A*β*40 and A*β*42 to induce platelet aggregation. The study of the role of platelets and the pathogenesis of Alzheimer’s disease is an active topic (Catricala *and others*, 2012).

We note that approximately half of genes in the high confidence gene set from the control vs. AD comparison (4282 genes) are also found in the high confidence gene sets from the control vs. PSP comparison (4207 genes). Altogether, 2158 DV genes are common to both comparisons. This observation is consistent with recent findings that transcriptomic changes are in AD and PSP relative to control are strongly correlated (Wang *and others*, 2022).

## 5. Discussion

In the present work, we have shown that using the CoDA framework for gene expression analysis leads to the natural modeling of CLR-transformed data using the skew-normal distribution. The latter is a tractable model with mature computational support through the `sn` R package. A test of differential variability can therefore be based directly on the standard deviation parameter of the skew-normal distribution. Moreover, a test of differential expression that is based on the mean parameter can be derived as well. With these tests, it becomes possible to develop methods for detecting three classes of genes in two-population comparisons: (i) equal variance, different mean; (ii) equal mean, different variance; (iii) different mean, different variance. Although clrDV cannot differentiate genes of the second and the third type, inspection of violin plots should be useful for ascertaining whether the DV genes also appear to differ significantly in the mean of their relative expression level.

For the control vs. AD and the control vs. PSP comparisons, we note that the high confidence gene set for which the estimated log_2_(SD ratio) has relatively large magnitude, already constitutes a substantial percentage (between 86.6% and 92.5%) of all DV genes called by clrDV. Thus, it seems that clrDV alone should be able to recover most of the DV genes of interest.

The relative poorer performance of MDSeq and GAMLSS could be caused by the choice of normalization. It is known that incorrect normalization leads to inflated FDR in differential expression analyses (Evans *and others*, 2018), yet the assumptions that justify a normalization method are usually not testable. Since existing normalization methods have been developed for the purpose of finding differentially expressed genes, the assumptions that justify their use are probably suboptimal for differential variability tests. Consequently, the performance of existing count-based approaches for DV test is likely sensitive to the choice of normalization method. However, it is beyond the scope of the present work to optimize the choice of normalization step for these count-based methods.

On the aspect of practical application, we note that the R codes provided by de Jong *and others* (2019) for GAMLSS are not sufficiently generic and require further user modifications to be suitable for routine use as a DV test. We also found that MDSeq may occasionally encounter difficulties in estimating model parameters. In the analysis of the Mayo RNA-Seq dataset, we found 14 and 11 genes returned NA parameter estimates in the analysis of the control vs. AD and control vs. PSP comparisons, respectively. Given these findings, we believe clrDV is currently the most practical and effective method for researchers who wish to conduct differential variability test using RNA-Seq data.

## 6. Code Availability

We have created an R package called `clrDV` to perform the differential variability test described here. The R package and codes for reproducing the analyses in this study are available at https://github.com/Divo-Lee/clrDV.

## 7. Supplementary Material

Supplementary material is available online at http://biostatistics.oxfordjournals.org.

## Acknowledgments

We wish to thank Dr. C.Y. Ung for helpful discussions. The results published here are in whole or in part based on data obtained from the AD Knowledge Portal (https://adknowledgeportal.org). The Mayo RNAseq study data was led by Dr. Nilüfer Ertekin-Taner, Mayo Clinic, Jacksonville, FL as part of the multi-PI U01 AG046139 (MPIs Golde, Ertekin-Taner, Younkin, Price). Samples were provided from the following sources: The Mayo Clinic Brain Bank. Data collection was supported through funding by NIA grants P50 AG016574, R01 AG032990, U01 AG046139, R01 AG018023, U01 AG006576, U01 AG006786, R01 AG025711, R01 AG017216, R01 AG003949, NINDS grant R01 NS080820, CurePSP Foundation, and support from Mayo Foundation. Study data includes samples collected through the Sun Health Research Institute Brain and Body Donation Program of Sun City, Arizona. The Brain and Body Donation Program is supported by the National Institute of Neurological Disorders and Stroke (U24 NS072026 National Brain and Tissue Resource for Parkinsons Disease and Related Disorders), the National Institute on Aging (P30 AG19610 Arizona Alzheimers Disease Core Center), the Arizona Department of Health Services (contract 211002, Arizona Alzheimers Research Center), the Arizona Biomedical Research Commission (contracts 4001, 0011, 05-901 and 1001 to the Arizona Parkinson’s Disease Consortium) and the Michael J. Fox Foundation for Parkinsons Research.

## Conflict of Interest

None declared.

## Supplementary Material for

### S1. The Skew-Normal Distribution

Let *ϕ*(*z*) be the standard normal probability density function 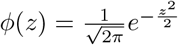, with cumulative distribution function 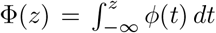. The probability density function of the skew-normal distribution is given by*ϕ*(*z, α*) = 2*ϕ*(*z*)Φ(*αz*), where the skewness parameter *α* ∈ ℝ. Suppose *Z* is a random variable that has the skew-normal distribution (i.e. *Z* ∼ SN(*α*)). Its mean and its variance are given by 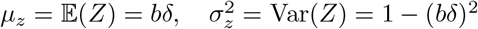, respectively, where 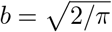 and 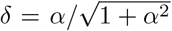. The formal derivation of the properties of the skew-normal distribution is due to Azzalini, 1985, who treated the skew-normal distribution as a generalization of the normal distribution. Historically, the skew-normal model were arrived at by several different authors in other contexts (e.g. as a prior distribution in Bayesian analysis by O’Hagan and Leonard, 1976; see Azzalini, 2022). However, these authors did not elaborate further on the theoretical properties of the skew normal as in Azzalini, 1985.

Figure S1 shows some examples of the probability density function of the skew-normal distribution with centered parameters for different parameter combinations.

#### S1.1 Fisher Information Matrix

For the direct parameters (DP) vector ***θ***^(*D*)^ = (*ξ, ω, α*), the Fisher information matrix is given by

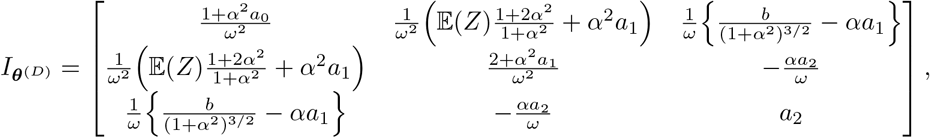

where 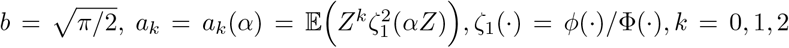 (Azzalini, 1985). The matrix 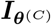 becomes singular as *α* → 0. This problem prevents the direct application of maximum likelihood estimation (MLE) for estimating the parameters of the DP form of the skew-normal distribution. In the same paper, Azzalini (1985) introduced the centered parametrization form to address the singulariy problem. He redefines a skew-normal variable

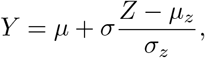

which has 𝔼(*Y*) = *µ* and Var(*Y*) = *σ*^2^. The centered parameters (CP) vector ***θ***^(*C*)^ = (*µ, σ, γ*) has parameter space ℝ × ℝ^+^ × (−0.9953, 0.9953) (Azzalini and Capitanio, 2014). The skewness parameter *γ* is the coefficient of skewness of *Z*, and also that of *Y*. Thus, we write *Y* ∼ SN_C_(*µ, σ, γ*). Fig. S1 shows examples of the probability density functions of the skew-normal distribution with CP for different values of *µ, σ* and *γ*.

For centered parameters ***θ***^(*C*)^ = (*µ, σ, γ*), the Fisher information matrix is given by

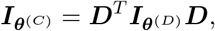

where ***D*** is the Jacobian matrix, that is, the derivatives of the parameters ***θ***^(*D*)^ = (*ξ, ω, α*) with respect to ***θ***^(*C*)^ = (*µ, σ, γ*). The Jacobian matrix ***D*** is given by

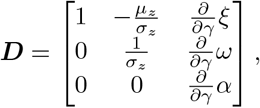

where *μ*_*z*_ = *bδ* and 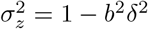. The terms in the last column of ***D*** are given by

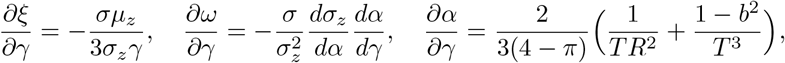

where

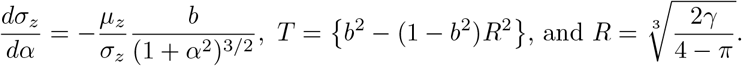

After some algebra, 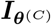 can be expressed as

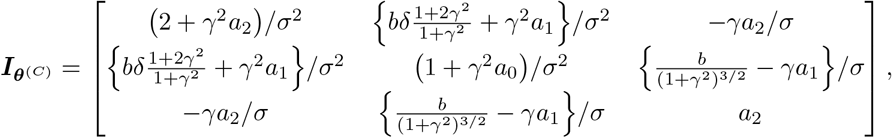

where

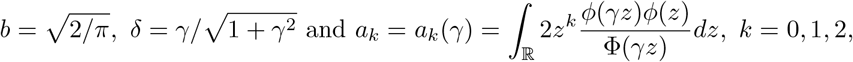

following the notation in Liseo and Loperfido (2006). The Fisher information matrix 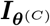 converges to a diagonal matrix with diagonal (1*/σ*^2^, 2*/σ*^2^, 1*/*6), as *γ* → 0 (Azzalini 1985; Azzalini and Capitanio, 2014).

## S2. List of R Packages Used

The following R packages (according to alphabetical order) were used in the present work: `compositions` (van den Boogaart *and others*, 2022), `edgeR` (Robinson *and others*, 2010), `gamlss` (Rigby and Stasinopoulos, 2005), `gridExtra` (Baptiste, 2017), `httr` (Wickham, 2022), `jsonlite` (Ooms, 2014), `MDSeq` (Ran and Daye, 2017), `missMethyl` (Phipson and Oshlack, 2014), `polyester` (Alyssa *and others*, 2022), `readr` (Hadley *and others*, 2022), `sn` (Azzalini, 2022), `VennDiagram` (Chen, 2022) and `vioplot` (Adler and Kelly, 2020).

## S3. Captions for Supplementary Tables

Supplementary Table 1: List of DV genes detected by clrDV for the control vs. AD comparison in the analysis of the Mayo RNA-Seq dataset.

Supplementary Table 2: List of DV genes detected by clrDV for the control vs. PSP comparison in the analysis of the Mayo RNA-Seq dataset.

Supplementary Table 3: List of DV genes detected by clrDV, MDSeq, and GAMLSS for the control vs. AD comparison in the analysis of the Mayo RNA-Seq dataset.

Supplementary Table 4: List of DV genes detected by clrDV, MDSeq, and GAMLSS for the control vs. PSP comparison in the analysis of the Mayo RNA-Seq dataset.

---

**Fig. S1.**
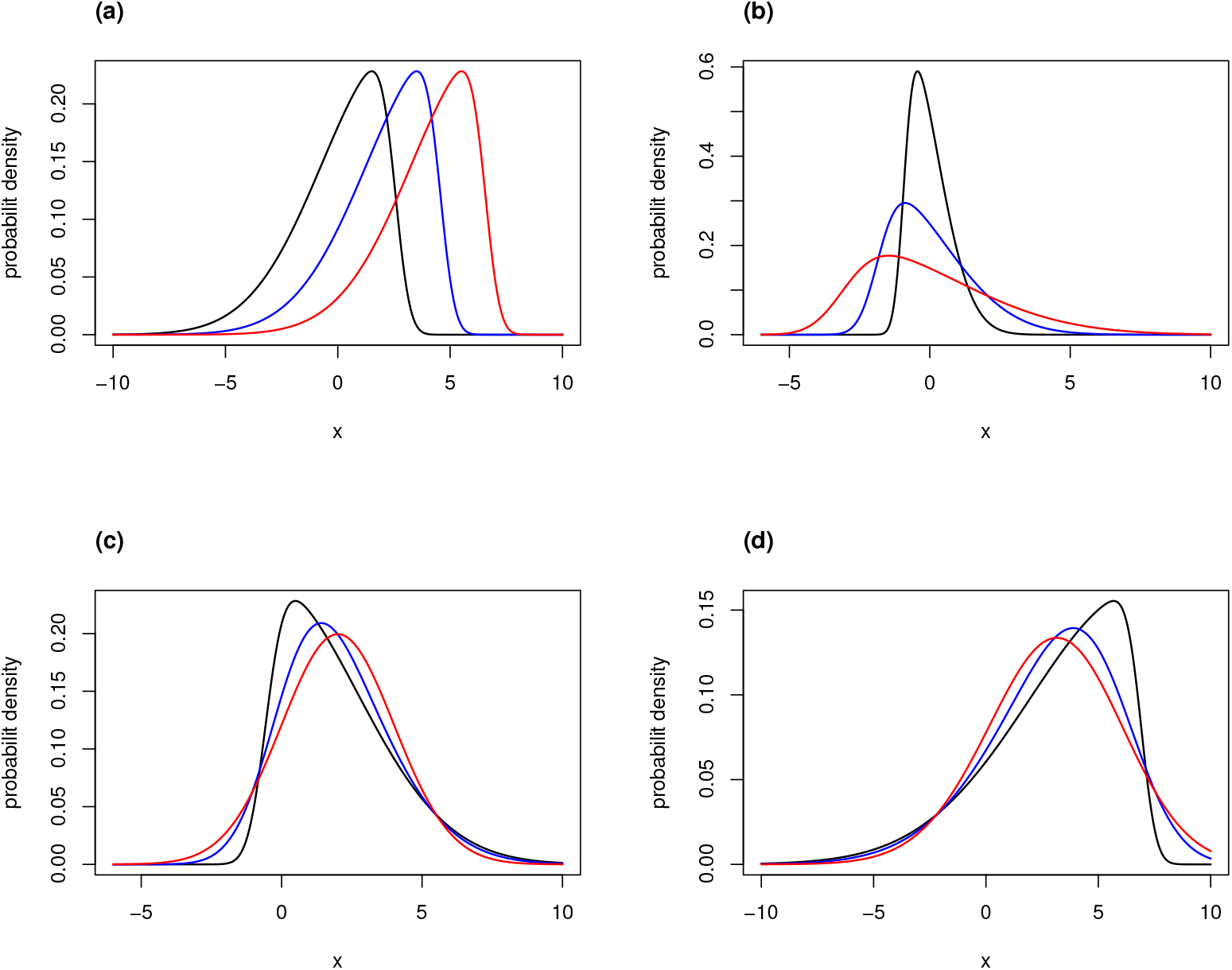
Probability density functions of the skew-normal distribution with centered parameters. (a) *σ* = 2, *γ* = *−*0.9; black: *µ* = 0, blue: *µ* = 2, red: *µ* = 4; (b) *µ* = 0, *γ* = 0.8; black: *σ* = 0.75, blue: *σ* = 1.5, red: *σ* = 2.5; (c) *µ* = 2, *σ* = 2; black: *γ* = 0.9, blue: *γ* = 0.5, red: *γ* = 0; and (d) *µ* = 3, *σ* = 3; black: *γ* = *−*0.95, blue: *γ* = *−*0.5, red: *γ* = *−*0.1.

## Notes

### Competing Interest Statement

The authors have declared no competing interest.

